# Discovery and Genomic Characterization of a Novel Henipavirus, Angavokely virus, from fruit bats in Madagascar

**DOI:** 10.1101/2022.06.12.495793

**Authors:** Sharline Madera, Amy Kistler, Hafaliana C. Ranaivoson, Vida Ahyong, Angelo Andrianiaina, Santino Andry, Vololoniaina Raharinosy, Tsiry H. Randriambolamanantsoa, Ny Anjara Fifi Ravelomanantsoa, Cristina M. Tato, Joseph L. DeRisi, Hector C. Aguilar, Vincent Lacoste, Philippe Dussart, Jean-Michel Heraud, Cara E. Brook

## Abstract

The genus *Henipavirus* (family *Paramyxoviridae*) is currently comprised of seven viruses, four of which have demonstrated prior evidence of zoonotic capacity. These include the biosafety level 4 agents Hendra (HeV) and Nipah (NiV) viruses, which circulate naturally in pteropodid fruit bats. Here, we describe and characterize Angavokely virus (AngV), a divergent henipavirus identified in urine samples from wild, Madagascar fruit bats. We report the near-complete 16,740 nt genome of AngV, which encodes the six major henipavirus structural proteins (nucleocapsid, phosphoprotein, matrix, fusion, glycoprotein, and L polymerase). Within the phosphoprotein (P) gene, we identify an alternative start codon encoding the AngV C protein and a putative mRNA editing site where the insertion of one or two guanine residues encodes, respectively, additional V and W proteins. In other paramyxovirus systems, C, V, and W are accessory proteins involved in antagonism of host immune responses during infection. Phylogenetic analysis suggests that AngV is ancestral to all four previously described bat henipaviruses—HeV, NiV, Cedar virus (CedV), and Ghanaian bat virus (GhV)—but evolved more recently than rodent- and shrew-derived henipaviruses, Mojiang (MojV), Gamak (GAKV), and Daeryong (DARV) viruses. Predictive structure-based alignments suggest that AngV is unlikely to bind ephrin receptors, which mediate cell entry for all other known bat henipaviruses. Identification of the AngV receptor is needed to clarify the virus’s potential host range. The presence of V and W proteins in the AngV genome suggest that the virus could be pathogenic following zoonotic spillover.

**Importance:** Henipaviruses include highly pathogenic emerging zoonotic viruses, derived from bat, rodent, and shrew reservoirs. Bat-borne Hendra (HeV) and Nipah (NiV) are the most well-known henipaviruses, for which no effective antivirals or vaccines for humans have been described. Here we report the discovery and characterization of a novel henipavirus, Angavokely virus (AngV), isolated from wild fruit bats in Madagascar. Genomic characterization of AngV reveals all major features associated with pathogenicity in other henipaviruses, suggesting that AngV could be pathogenic following spillover to human hosts. Our work suggests that AngV is an ancestral bat henipavirus which likely uses viral entry pathways distinct from those previously described for HeV and NiV. In Madagascar, bats are consumed as a source of human food, presenting opportunities for cross-species transmission. Characterization of novel henipaviruses and documentation of their pathogenic and zoonotic potential are essential to predicting and preventing the emergence of future zoonoses that cause pandemics.

## Introduction

Henipaviruses (HNVs) belong to a genus of bat-, rodent-, and shrew-borne viruses within the family *Paramyxoviridae* with demonstrated zoonotic potential. HNVs can manifest extreme virulence in human hosts, as exemplified by the prototypical HNVs, Hendra virus (HeV), and Nipah virus (NiV), which cause severe acute respiratory distress and/or encephalitis in humans, yielding case fatality rates that can exceed 90% (1–3). This high pathogenicity and the lack of approved HNV therapeutics or vaccines for humans have garnered HeV and NiV classification as Biological Safety Level 4 (BSL4) agents and WHO priority diseases. Since their discovery in the 1990s, HeV and NiV have periodically spilled over to humans from their reservoir hosts, pteropodid bats. HeV zoonosis is mediated by spillover to intermediate horse hosts, from which humans acquire infection(4). NiV can spillover to humans via intermediate transmission through pig hosts, or directly from bat-to-human, resulting in near-annual outbreaks of fatal encephalitis in South Asia, where subsequent human-to-human transmission also occurs (2, 5– 7).

Novel HNVs continue to emerge from wildlife hosts and represent ongoing threats to human health. Initially, the *Henipavirus* genus comprised only HeV and NiV; however, the past two decades have witnessed the discovery of five new HNVs: bat-borne Cedar virus (CedV) and Ghanaian bat virus (GhV), rodent-borne Mojiang virus (MojV), and shrew-borne Gamak (GAKV) and Daeryong viruses (DARV) (8–11). Of these novel HNVs, at least two show evidence of zoonotic potential: serological data suggests prior human exposure to GhV in West Africa (12), while MojV was first identified following a human outbreak of severe pneumonia in Chinese mine workers, all of whom died after infection(9). In addition to their high potential for pathogenicity, HNVs possess a broad host range that spans at least seven mammalian orders, including bats (10, 13).

Cross-species viral spillover necessitates effective inter-species transmission, which first requires a virus to successfully enter the cells of diverse host species. In general, HNVs use the highly-conserved ephrin family of proteins, both type A and type B, as cell entry receptors (1, 8, 14–16). A notable exception to this pattern is MojV, which does not use ephrin proteins—or the sialic acid and CD150 receptors common to non-HNV paramyxoviruses—to gain cell entry (14, 17). Indeed, as of yet, the viral entry receptor for MojV—and the closely related GAKV and DARV—remain unknown. In general, viruses in the genus *Henipavirus* have broad host ranges and cause high case fatality rates following human spillover, making the characterization of new HNVs a high public health priority.

The HNV genome consists of six structural proteins: nucleocapsid (N), phosphoprotein (P), matrix (M), fusion (F), glycoprotein (G), and polymerase (L). In comparison with other members of the family *Paramyxoviridae*, HNVs have relatively larger genomes (approximately 18kb vs 16kb). This extended length is largely due to several, long 3’ untranslated regions (UTR) of the N, P, F and G mRNAs (18, 19). The genome length of HNVs, like all paramyxoviruses, adheres to the so-called ‘Rule of Six’, whereby viral genomes consistently demonstrate polyhexameric length (20). The ‘Rule of Six’ is believed to be a requirement for efficient genome replication under the unique mRNA editing features of the paramyxovirus genome (20). The paramyxovirus P locus exhibits notable transcription properties that are shared across most members of the *Paramyxoviridae* family. The P gene permits the translation of additional accessory proteins from either gene editing events within the locus (prior to translation) or an overlapping ORF in the P gene itself. All HNVs, with the exception of CedV, harbor a highly conserved mRNA editing site at which the insertion of additional guanine residues can result in the translation of accessory proteins, V and W, involved in viral antagonism and evasion of the host immune system (1). The HNV P gene also contains an overlapping ORF that allows for the synthesis of a third accessory protein, C, which is also involved in viral host immune evasion (1).

Our lab has previously demonstrated evidence of exposure to henipa-like viruses in serum collected from three endemic Madagascar fruit bat species (*E. dupreanum, Pteropus rufus*, and *Rousettus madagascariensis*) using a Luminex serological assay which identified cross reactivity to CedV/NiV/HeV-G and -F proteins (21). The most significant antibody binding previously detected corresponded to the NiV-G antigen for *E. dupreanum* serum and the HeV-F antigen for *P. rufus* and *R. madagascariensis* serum, suggesting the potential circulation of multiple HNVs in the Madagascar fruit bat system (21). Fruit bats, including *E. dupreanum*, are consumed widely in Madagascar as a source of human food, presenting opportunities for cross-species zoonotic emergence. This underscores the importance of further characterization of the pathogenic and zoonotic potential of AngV and other potential HNVs circulating in the Madagascar fruit bat system. Here, we describe and characterize a novel bat HNV, Angavokely virus (AngV), recovered from urine samples collected from the Madagascar fruit bat, *E. dupreanum*. Our work suggests AngV is part of an ancestral group of HNVs that may rely on a novel, non-ephrin-mediated viral entry pathway.

## Methods

### Ethics Statement

Animal capture and handling and subsequent collection of biological samples were conducted in strict accordance with the Madagascar Ministry of Forest and the Environment (permit numbers 019/18, 170/18, 007/19) and guidelines posted by the American Veterinary Medical Association. Field protocols were approved by the UC Berkeley Animal Care and Use Committee (ACUC Protocol # AUP-2017-10-10393), as previously described (22).

### Animal capture, sample collection, and RNA extraction

Fruit bats were captured and processed in part with a long-term study investigating the seasonal dynamics of potentially zoonotic viruses in Madagascar, as has been previously described (21–25). Animals were identified morphologically by species, sex, and age class (juvenile vs. adult), and urine swabs were collected into viral transport medium from any individual that urinated during handling. Urine swabs were flash-frozen in liquid nitrogen in the field and delivered to -80^0^C freezers at Institut Pasteur de Madagascar for long-term storage. Urine specimens from 206 bats were randomly selected for total RNA extraction using the Zymo Quick RNA/DNA Microprep Plus kit, performed as previously described (22).

### mNGS library preparation

Total urine RNAs were diluted with nuclease-free H_2_O, and 5uL of each specimen was used as input for mNGS library preparation. A 2-fold dilution series of a 25ng/uL stock of HeLa total RNA (n=8 samples), along with 5 water samples were included and processed in parallel as positive and negative controls, respectively. Additionally, a 25pg aliquot of External RNA Control Consortium (ERCC) spike-in mix (Thermo-Fisher) was included in each sample. Dual-indexed mNGS library preparations for the samples were miniaturized and performed in 384-well format with NEBNext Ultra II RNAseq library preparation kit (New England Biolabs) reagents. RNA samples were fragmented for 12 min at 94°C, and 16 cycles of PCR amplification were performed. Per sample read yields from a small scale iSeq (Illumina) paired-end 2 × 146bp sequencing run on an equivolume pool of the individual libraries were used to normalize volumes of the individual mNGS libraries to generate an equimolar pool. Paired-end 2 × 146bp sequencing of the resulting equimolar library pool was performed on the NovaSeq6000 (Illumina) to obtain approximately 50 million reads per sample.

### Sequence analysis

Raw reads from urine sample sequencing were first uploaded to the CZBID (v6.8) platform for host and quality filtering and *de novo* assembly (26). In brief, in the CZBID pipeline, adaptor sequences were removed with Trimmomatic (v.0.38), and reads were quality filtered (27). Reads then underwent host-filtration against the Malagasy fruit bat genome, *E. dupreanum*, using STAR (v 2.7.9a) (28) and a second host-removal step using Bowtie2 (29). After host filtering, reads were aligned using rapsearch2 (30) and Gsnap (31), and putative pathogen taxa were identified. Next, reads were assembled using SPADES (v.3.15.3) (32), and all contigs generated were subject to BLAST analysis against the putative taxa previously identified by rapsearch2 and Gsnap. We considered samples positive for HNV if the CZBID pipeline produced at least one contig with an average read depth of two or more, which yielded a BLAST alignment length >100 nt/aa and an e-value < 0.00001 (BLASTn v2.5.0+) or a bit score >100 (BLASTx v2.5.0+) when queried against an HNV database derived from all HNV genomes available in NCBI (Accessed July 2021).

### Genome Annotation and Comparison

One urine sample, collected from an adult female *E. dupreanum* fruit bat in March 2019, yielded a near full-length HNV genome, which we analyzed in greater depth in subsequent analyses and annotated as the novel HNV, AngV (Genbank Accession #: ON613535). Nucleotide BLAST of the AngV genome identified NiV (GenBank Accession #: AF212302) as the top hit for this novel virus and was subsequently chosen as the reference genome for further analysis. We aligned AngV to NiV (GenBank Accession #: AF212302) in the program Geneious Prime (v2020.2.4) and annotated all six major HNV structural genes, and the accessory C ORF, within the P gene. We identified the putative mRNA editing site within the P gene sequence (spanning nucleotides 1,225-1,232 of the P gene) and manually added one or two guanine (G) residues to the 3’ end of the conserved HNV mRNA editing site to generate V and W ORFs, respectively, and their corresponding proteins. We furthered queried all identified transcriptional elements against publicly available sequences using NCBI BLAST and BLASTx (33). Resulting BLAST and BLASTx hits were used in phylogenetic analyses as described below.

We used the program pySimPlot to scan the whole genome sequence of AngV for nucleotide sequence identity to the NiV genome (GenBank Accession #: AF212302) and the nucleotide and amino acid sequences of individual Open Reading Frames (ORFs) contained therein. Respectively, window size and scanning were specified as 50 and one for nucleotide pairwise identity and 50 and five for amino acid pairwise identity. Results were visualized using Prism (9.2.0).

### Phylogenetic analyses

We constructed 10 Maximum Likelihood (ML) phylogenetic trees to analyze the evolutionary relatedness of our putative HNV to previously described paramyxoviruses. These included (a) one amino acid L protein phylogeny comparing the L protein of AngV to all reference L protein paramyxovirus sequences in NCBI, in addition to the newly-described shrew HNV, GAKV and DARV (accessed November 2021) and (b) nine amino acid phylogenies comparing each individual protein annotated in AngV (N, P, C, V, W, M, F, G, L) against the top 50 BLASTx sequence hits for each protein collapsed on 98% sequence similarity. Distinct outgroups were applied: (a) Sunshine Coast Virus (GenBank Accession #: YP_009094051.1) for L protein, (b) Human orthopneumovirus (HRSV, GenBank Accession #: NC_001781) for N, P, M, F, G, L proteins, and (c) Sendai virus (GenBank Accession #: NP_056872) for gene C.

For each phylogenetic tree, we aligned sequences via the MUSCLE algorithm (v3.8.1551) (34) and determined the best fit nucleotide or amino acid substitution model for using ModelTest-NG (35). Phylogenies were then constructed in RAxML-NG (36), using the corresponding best fit model: JTT (complete L-protein sequence) or LG+G4+F (individual proteins). In accordance with best practices outlined in the RAxML-NG manual, twenty ML inferences were made on each original alignment. Bootstrap replicate trees were inferred using Felsenstein’s method (37). MRE-based bootstopping test was applied after every 200 replicates (38), and bootstrapping was terminated once diagnostic statistics dropped below the threshold value. Bootstrap support values were drawn on the best-scoring tree.

We additionally computed one Bayesian time-resolved phylogeny, using all 77 full-length HNV nucleotide sequences available in NCBI, including our newly contributed AngV (GenBank Accession #: ON613535). As with ML trees, sequences were first aligned in MUSCLE (v3.8.1551) (34), and the best fit nucleotide substitution model was subsequently queried in ModelTest-NG (35). We then constructed a Bayesian timetree in the program BEAST2 (39, 40), using the best fit GTR+I+G4 model inferred for the whole genome alignment from ModelTest-NG and assuming a constant population prior. Sampling dates corresponded to collection data as reported in NCBI Virus; we assumed a collection date of 31-July in cases where only year of collection was reported. We computed trees using both an uncorrelated exponentially distributed relaxed molecular clock (UCED) and a strict clock but here report results from the strict clock only as similar results were inferred from both. We ran Markov Chain Monte Carlo (MCMC) sample chains for 1 billion iterations, checked convergence using TRACER v1.7 (41) and averaged trees after 10% burn-in using TreeAnnotator v2.6.3 (42) to visualize mean posterior densities at each node. The resulting phylogeny was visualized in R v.4.0.3 for MacIntosh in the ‘ggtree’ package (43).

### AngV G Protein Structure Modeling

We used the Artificial Intelligence system, AlphaFold, to predict the 3D structure of the AngV glycoprotein (G) (44). Molecular graphics and analyses of the AngV glycoprotein structure were performed with UCSF ChimeraX (45). HNV glycoprotein ephrin binding residues were aligned using the program Geneious Prime (v2020.2.4).

## Results

### Discovery and prevalence of HNV in Malagasy bats

Urine swab specimens from 206 bats were collected in 8 roosting sites across the island of Madagascar from 2013 to 2019 (Figure 1). Urine samples were collected during wet and dry seasons from all three Madagascar fruit bat species, *P. rufus, E. dupreanum*, and *R. madagascariensis* (Table 1). Isolated RNA from urine swab specimens generated an average of 19 million paired-read sequences. In total, 10/206 (4.9%) bats were positive for HNV; all positive samples were collected from *E. dupreanum* bats (10/106; 9.4%) at the Angavokely cave roosting site (Table 1). Positive samples were collected in wet and dry seasons from both male and female adults.

**Figure 1.**
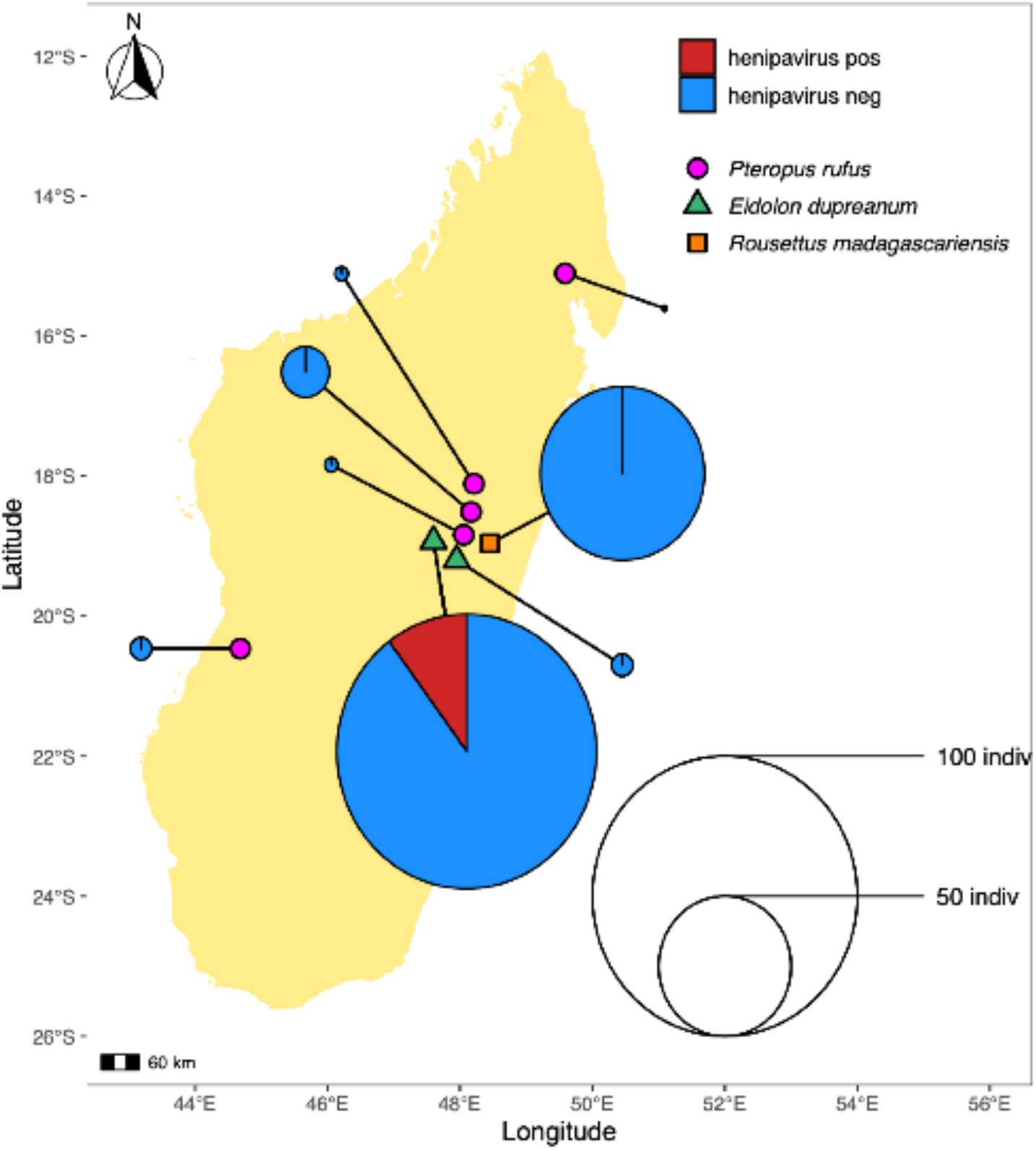
Geographic location of sampling sites used in this study. Sampling sites grouped by bat species found depicted as follows *P. rufus* (pink circles) Ambakoana (−18.51 S, 48.17 E) / Mahabo (−20.46 S, 44.68 E) / Mahialambo (−18.11 S, 48.21 E) / Makira (−15.11 S, 49.59 E) / Marovitsika (−18.84 S, 48.06 E) roosts; *E. dupreanum* (green triangles) Angavobe (−18.94 S, 47.95 E) /Angavokely (−18.93 S, 47.76 E) caves; *R. madagascariensis* (orange squares) Maromizaha cave (−18.96 S, 48.45). Pie charts are size-weighted by total bat population sampled at each site, corresponding to the legend. The percentage of HNV positive samples is shown for all sampled species and sites. HNV positive samples were only recovered from the *E. dupreanum* Angavokely site.

**Table 1.**
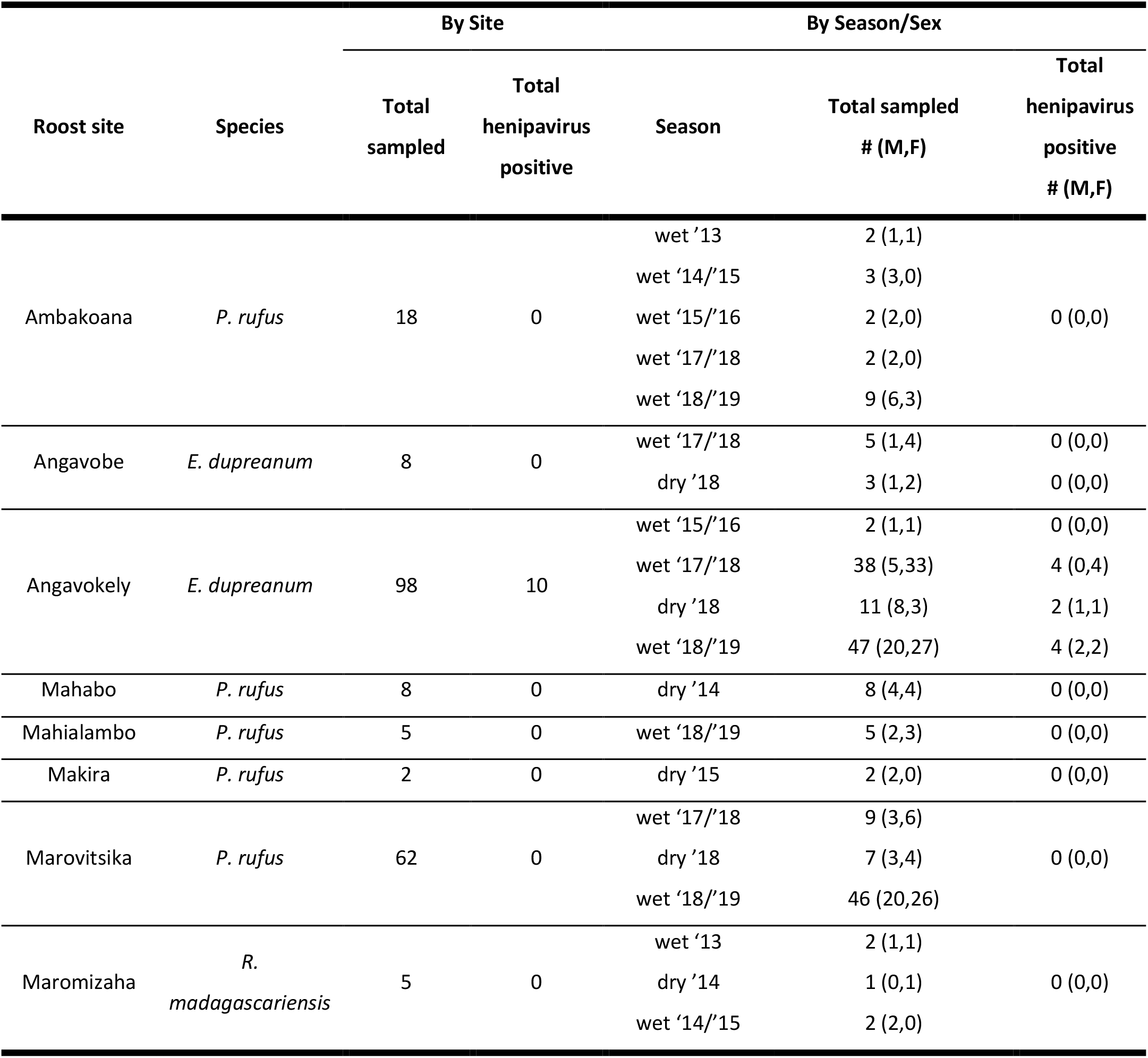
Prevalence of HNV infections in the urine of Malagasy bats captured during 2013-2019 collection period.

| Roost site | Species | By Site |  | By Season/Sex |  |  |
| --- | --- | --- | --- | --- | --- | --- |
|  |  | Total sampled | Total henipavirus positive | Season | Total sampled # (M,F) | Total henipavirus positive # (M,F) |
| Ambakoana | *P. rufus* | 18 | 0 | wet '13 | 2 (1,1) | 0 (0,0) |
|  |  |  |  | wet '14/'15 | 3 (3,0) |  |
|  |  |  |  | wet '15/'16 | 2 (2,0) |  |
|  |  |  |  | wet '17/'18 | 2 (2,0) |  |
|  |  |  |  | wet '18/'19 | 9 (6,3) |  |
| Angavobe | *E. dupreanum* | 8 | 0 | wet '17/'18 | 5 (1,4) | 0 (0,0) |
|  |  |  |  | dry '18 | 3 (1,2) | 0 (0,0) |
| Angavokely | *E. dupreanum* | 98 | 10 | wet '15/'16 | 2 (1,1) | 0 (0,0) |
|  |  |  |  | wet '17/'18 | 38 (5,33) | 4 (0,4) |
|  |  |  |  | dry '18 | 11 (8,3) | 2 (1,1) |
|  |  |  |  | wet '18/'19 | 47 (20,27) | 4 (2,2) |
| Mahabo | *P. rufus* | 8 | 0 | dry '14 | 8 (4,4) | 0 (0,0) |
| Mahialambo | *P. rufus* | 5 | 0 | wet '18/'19 | 5 (2,3) | 0 (0,0) |
| Makira | *P. rufus* | 2 | 0 | dry '15 | 2 (2,0) | 0 (0,0) |
| Marovitsika | *P. rufus* | 62 | 0 | wet '17/'18 | 9 (3,6) | 0 (0,0) |
|  |  |  |  | dry '18 | 7 (3,4) |  |
|  |  |  |  | wet '18/'19 | 46 (20,26) |  |
| Maromizaha | *R. madagascariensis* | 5 | 0 | wet '13 | 2 (1,1) | 0 (0,0) |
|  |  |  |  | dry '14 | 1 (0,1) |  |
|  |  |  |  | wet '14/'15 | 2 (2,0) |  |

### Genomic characterization of AngV

We recovered one near-full-length HNV contig (16,740 nt), supported by an average sequencing depth of 14 reads (Figure 2A), from a urine sample collected from an adult, non-lactating *E. dupreanum* female in the 2018-2019 wet season (capture date: 15-March 2019). We focused subsequent genomic analyses on this longest sequence, which we named Angavokely virus (AngV) after the site of *E. dupreanum* capture.

**Figure 2.**
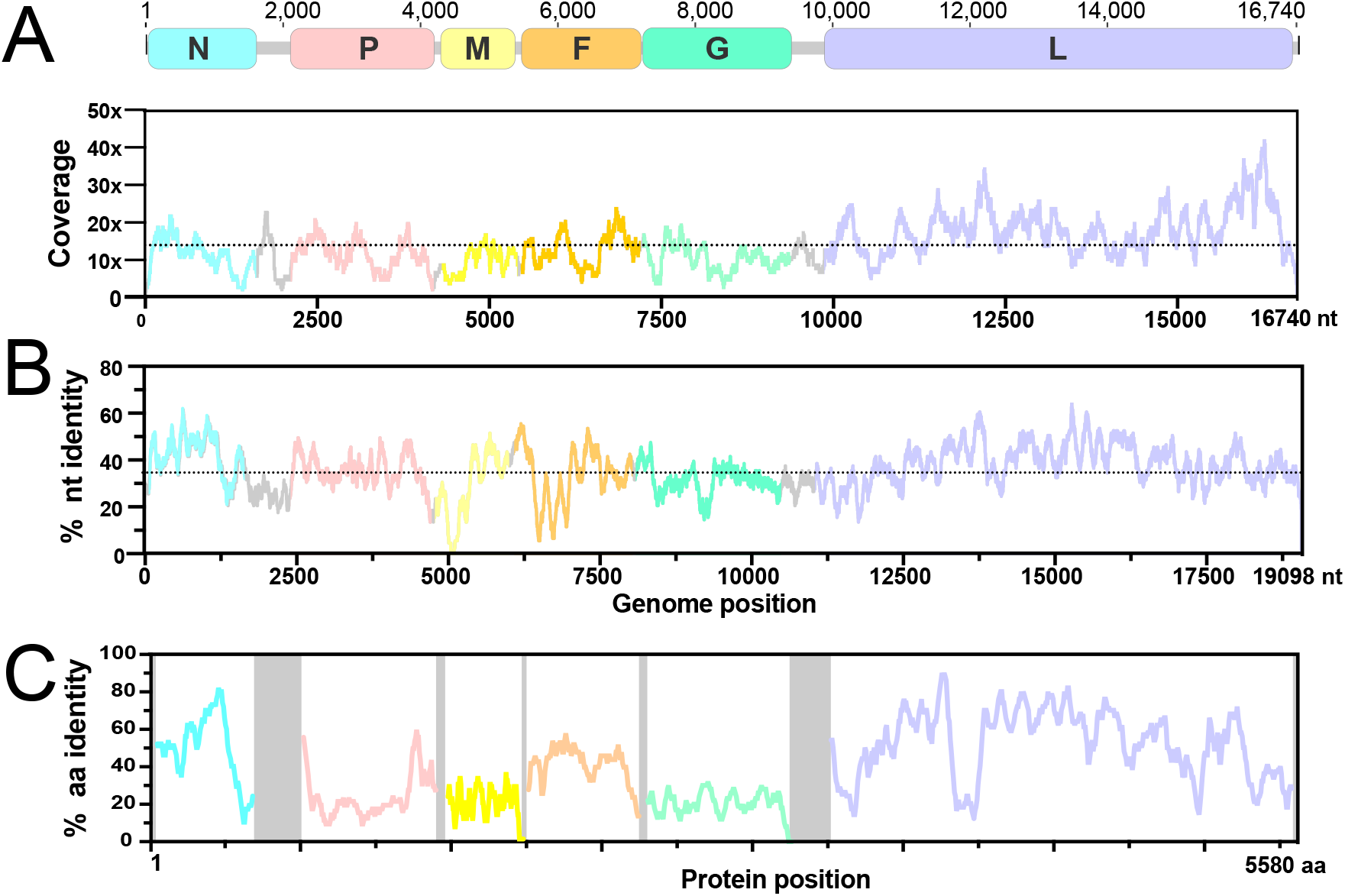
AngV genome organization. A. Coding regions for each gene are shown and depicted in color, non-coding intergenic and terminal regions are highlighted in gray. Depicted genes represented as follows: nucleocapsid (N), RNA polymerase (P and L), matrix (M), fusion (F), and glycoprotein (G). Sequencing read depth supporting each position of the recovered genome sequence is plotted below the genomic schematic. Scanning nucleotide (B) and amino acid (C) pairwise identity to Nipah virus (GenBank Accession #: AF212302). Dotted horizontal lines represent average read depth (14.29) or average nucleotide pairwise identity (36%).

As with other members of the *Henipavirus* genus, the genome of AngV is organized into 6 open reading frames (ORF) arranged in the order 3’-N-P-M-F-G-L-5’. AngV shares an average nucleotide identity of 36% with the NiV reference genome (AF212302) and a varying amino acid identity that is highest across the ORFs encoding for the nucleocapsid and L polymerase proteins (Figures 2B, 2C).

### AngV coding regions

The P gene of AngV follows an organization similar to most members of the *Henipavirus* genus. AngV harbors alternative start sites which, respectively, encode the P and C proteins, as well as a conserved putative mRNA editing site common to most paramyxoviruses A^4-6^G^2-3^ (Figure 3A and B). AngV shares an identical putative mRNA editing site with the recently discovered HNVs, MojV and GAKV. Pseudotemplated addition of one or two G residues at the conserved putative mRNA editing site generates a putative V and W protein, respectively (Figure 3C; W protein in Supplemental Figure 1). In congruence with members of the *Henipavirus* genus that encode the conserved putative mRNA editing site, the putative V protein of AngV harbors a unique C-terminal region that contains a highly conserved cysteine-rich zinc finger domain (Figure 3C).

**Figure 3.**
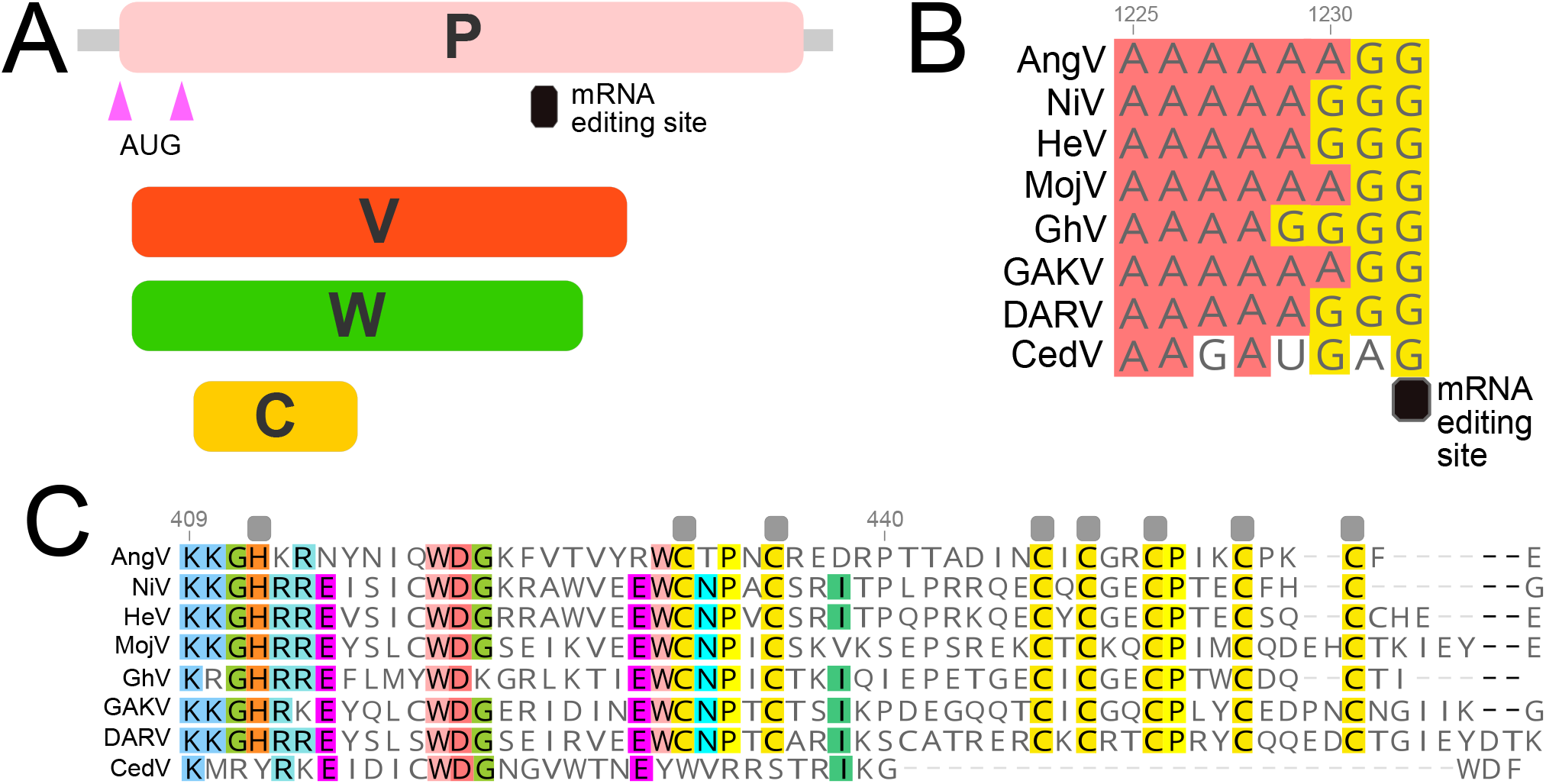
Organization of the P gene of AngV. A. Alternative transcriptional start sites (pink triangle) generate the P and C protein. Pseudotemplated addition of one or two guanine nucleotides at the putative mRNA editing site generates a V and W protein, respectively. B. Sequence alignment of the putative mRNA editing site across members of the *Henipavirus* genus (cRNA depicted). C. Amino acid alignment of the unique C terminal region of the V protein following the addition of one guanine nucleotide to the putative mRNA editing site. Gray boxes denote conserved cysteine and histidine residues suggested to directly coordinate bound zinc ions (54). Individual nucleotides or amino acids are color coordinated if at least 75% conserved at the alignment position. Nucleotide or amino acid position numbers displayed represent the position within the AngV gene or protein. Virus name (abbreviation), followed by GenBank Accession #: Angavokely virus (AngV) ON613535; Nipah virus (NiV) AF212302; Hendra virus (HeV) AF017149; Mojiang virus (MojV) KF278639; Ghanaian bat Henipavirus (GhV) HQ660129; Daeryong virus (DARV) MZ574409; Gamak virus (GAKV) MZ574407, Cedar virus (CedV) JQ001776. CedV is shown here only for comparison, as the CedV P protein is not believed to undergo RNA editing or to generate a functional V protein (8, 54).

The length of each ORF in the AngV genome resembles those from previously described HNVs, with the exception of the gene encoding the glycoprotein (G)—which, at 688 aa, is 56 aa longer than the longest previously-characterized HNV glycoprotein from GhV (632 aa; Table 2) (15). BLAST analysis indicates that AngV ORFs for genes encoding the nucleocapsid (N), matrix (M), and polymerase (L) proteins exhibit the highest nucleotide and amino acid pairwise identity with other HNVs, with highest similarity shared with the NiV L protein (nt 74.7%, aa 52.2%, Table 2). In contrast to many emerging viruses, AngV largely exhibits higher nucleotide vs. amino acid identity with other HNVs (Table 2). The more recently discovered HNVs (MojV, CedV, GAKV, DARV, GhV) mirror this pattern, showing higher nucleotide vs. amino acid identity when compared to NiV and HeV (data not shown).

**Table 2.** Length and pairwise sequence identity of predicted open reading frames of AngV and other HNV.

| Gene | AngV<br>length aa | NiV<br>length aa<br>(%nt, %aa) | HeV | MojV | CedV | GAKV | DARV | GhV |
| --- | --- | --- | --- | --- | --- | --- | --- | --- |
| N | 514 | 532<br>(56.4, 48.0) | 532<br>(55.4, 46.7) | 539<br>(55.8, 45.3) | 510<br>(57.5, 48.3) | 533<br>(58.0, 47.5) | 574<br>(55.3, 24.0) | 514<br>(56.2, 48.9) |
| P | 693 | 709<br>(47.8, 25.1) | 707<br>(48.1, 24.1) | 694<br>(47.6, 23.6) | 737<br>(42.3, 21.6) | 586<br>(43.8, 22.5) | 698<br>(47.3, 23.2) | 870<br>(42.2, 23.1) |
| C | 173 | 166<br>(51.3, 25.7) | 166<br>(51.5, 25.1) | 177<br>(47.6, 22.5) | 177<br>(54.5, 29.8) | 184<br>(47.4, 27.5) | 175<br>(48.1, 24.0) | 163<br>(46.1, 23.1) |
| V | 461 | 456<br>(46.9, 22.8) | 457<br>(48.5, 23.6) | 464<br>(47.2, 24.5) |  | 370<br>(43.0, 22.3) | 468<br>(47.2, 22.2) | 621<br>(24.7, 19.8) |
| W | 412 | 450<br>(47.0, 21.3) | 448<br>(45.0, 20.0) | 435<br>(46.5, 22.5) |  | 331<br>(43.9, 22.7) | 437<br>(45.5, 21.1) | 572<br>(39.3, 16.8) |
| M | 354 | 352<br>(60.0, 54.6) | 352<br>(59.5, 54.0) | 340<br>(57.1, 53.4) | 360<br>(56.9, 53.1) | 340<br>(56.1, 50.8) | 345<br>(56.5, 52.3) | 343<br>(56.9, 51.4) |
| F | 539 | 546<br>(53.7, 39.4) | 546<br>(51.7, 39.4) | 545<br>(52.2, 40.6) | 557<br>(51.8, 36.3) | 565<br>(53.3, 39.4) | 545<br>(58.8, 40.1) | 662<br>(45.5, 35.4) |
| G | 688 | 602<br>(43.5, 21.4) | 604<br>(43.8, 20.4) | 625<br>(43.0, 21.2) | 622<br>(43.3, 19.8) | 635<br>(44.9, 17.8) | 628<br>(44.3, 18.7) | 632<br>(42.5, 19.8) |
| L | 2,259 | 2,244<br>(74.7, 52.2) | 2,244<br>(57.6, 51.9) | 2,277<br>(57.5, 51.8) | 2,501<br>(52.4, 44.8) | 2,291<br>(57.7, 51.0) | 2,271<br>(58.8, 51.3) | 2,250<br>(57.0, 49.1) |

### AngV non-coding regions

Examination of all viral intergenic regions (in cRNA orientation) reveals that AngV exhibits the highly conserved CTT intergenic junction site characteristic of other HNVs, as well as gene stop and gene start sites with high similarity to those of previously described HNVs (Supplemental Table 1). We were unable to locate the intergenic junction site and transcriptional start or stop site in the 5’ region of the N ORF for AngV, suggesting that the genomic 3’ untranslated regions (UTR) for AngV have not yet been fully recovered. Comparison of the 5’ and 3’ UTRs for AngV with those of other HNVs reveals UTRs of varying lengths within the *Henipavirus* genus (Supplemental Table 2). Nevertheless, AngV exhibits similar lengths and a nucleotide identity of roughly 30-40% for the 5’ and 3’ UTRs of most HNV genes; however, the P gene 3’ UTR, the M gene 5’ and 3’ UTRs, and the F gene 5’ and 3’ UTRs, are significantly shorter in AngV compared with previously described HNVs. Correspondingly, nucleotide identity varies when comparing this shorter subset of 5’ and 3’ UTRs for AngV against other HNVs (Supplemental Table 2).

### Phylogenetic Analyses

Phylogenetic analysis of complete L protein amino acid sequences across the *Paramyxoviridae* family places AngV within the *Henipavirus* genus at <0.82 nucleotide substitutions away from the node distinguishing the family *Paramyxoviridae* from the *Sunviridae* (Figure 4A). AngV clusters independently within the *Henipavirus* genus and diverges ancestral to all currently known bat-borne HNVs. Our time-resolved Bayesian phylogeny further corroborates this result, placing AngV ancestral to all previously described bat-borne HNVs but more recently diverged than the rodent- and shrew-borne HNVs, MojV, GAKV, and DARV (Figure 4B; Supplemental Figure 2). We estimate the divergence of the AngV lineage from the rest of the HNV clade at 9,794 years ago (95% HPD 6,519-14,024 years), and the time to the Most Recent Common Ancestor (MRCA) for the entire HNV genus as 11,195 years ago (95% HPD 7,351-15,905 years).

**Figure 4.**
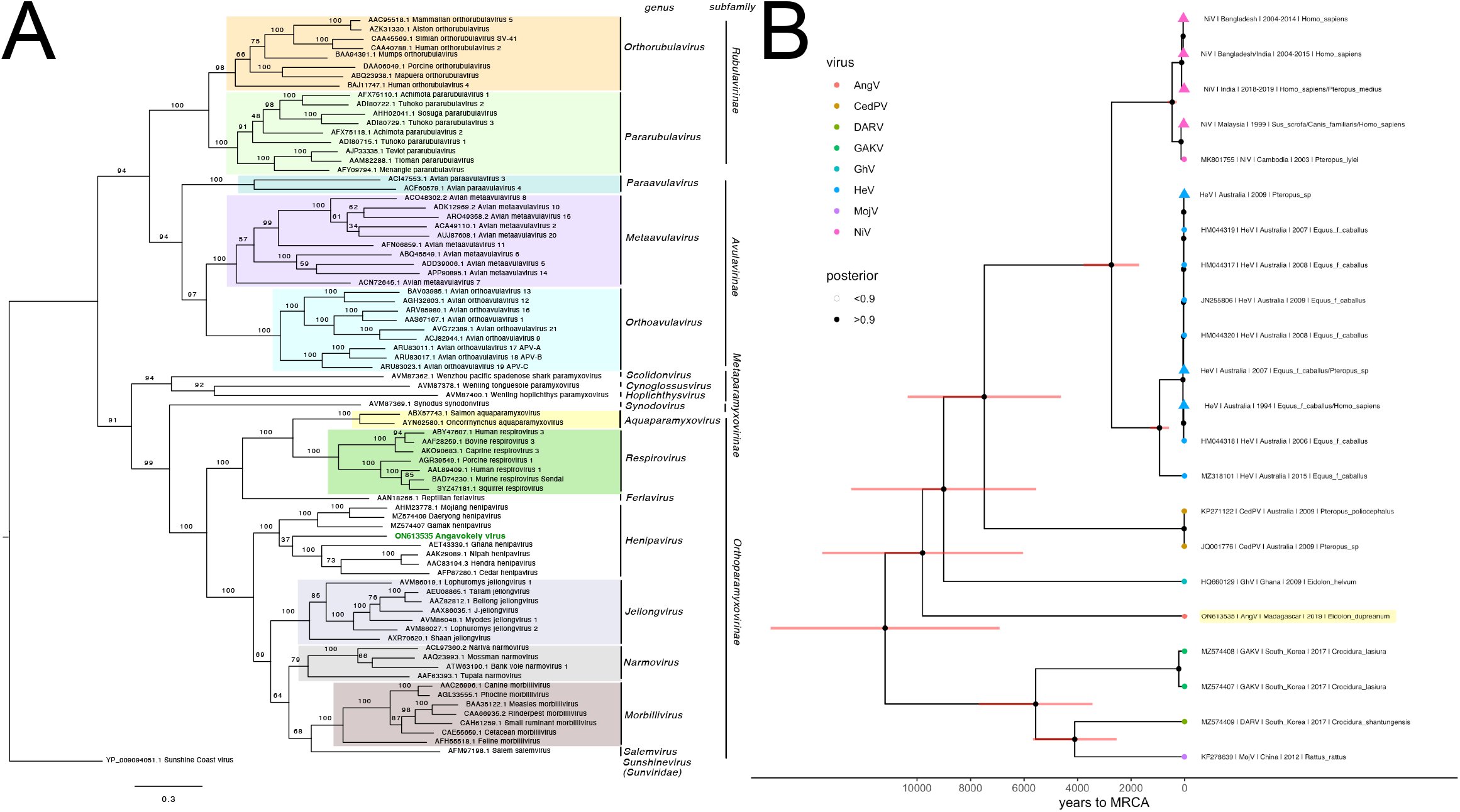
A. Phylogenetic analysis of the complete L protein sequences of members of the family *Paramyxoviridae*. Tree is rooted with Sunshine Coast Virus (GenBank Accession #: YP_009094051.1) as an outgroup, with outgroup branch length shrunk for ease of viewing. Novel HNV, AngV, is depicted in green. Subfamilies and genera are demarcated, excluding those unassigned to subfamily (genera *Scolidonvirus, Cynoglossusvirus, Hoplichthysvirus*). Bootstrap support is depicted and GenBank Accession numbers displayed next to virus names. Scale bar represents substitutions per site. B. Time-resolved Bayesian phylogeny computed in BEAST2 incorporating all available *Henipavirus* whole genome nucleotide sequences, with the addition of newly discovered GAKV, DARV, and AngV. Closely-related sequences are collapsed at triangle nodes for NiV and HeV (phylogeny with un-collapsed branches available in Supplemental Figure 2). 95% HPD intervals around the timing of each branching node are visualized as red horizontal bars. Posterior support >.9 is indicated by black coloring of the corresponding node, and distinct *Henipavirus* species are indicated by colored tip points, with AngV highlighted in yellow for further emphasis. The estimated time to MRCA for Angavokely virus and the previously-described bat-borne HNVs is 9,794 (95% HPD 6,519 – 14,025) years ago.

In N, P, C, V, W, M and G amino acid phylogenies, the AngV proteins cluster closely with those of other HNVs (Supplemental Figure 3). Interestingly, in the amino acid phylogeny, the AngV F protein, like the F proteins of MojV, GAKV, and DARV, localizes ancestral to non-HNV paramyxoviruses and distinct from the bat-borne HNV clade (Supplemental Figure 3). The AngV L protein shows the highest amino acid identity to the L protein of rodent-borne Mount Mabu Lophuromys virus 2 (MMLV-2), a putative Jeilongvirus (46), but is nonetheless nested between the MojV/GAKV/DARV clade and the bat-borne HNV clade (Supplementary Figure 3).

### AngV glycoprotein

We further examined the AngV G protein for conserved structural features and amino acid residues historically associated with HNV ephrin binding. AlphaFold analysis revealed a six-bladed β-propeller fold that is characteristic of *Paramyxoviridae* glycoproteins, with each blade largely composed of 4 antiparallel β-strands (Figure 5A). The β-propeller fold is stabilized by seven disulfide bonds that are conserved among HNVs (Supplemental Figure 4). This HNV protein G structure-based alignment reveals that the elongated AngV G protein primarily results from a lengthy C-terminal tail with an additional 67 aa beyond that of NiV G protein (Supplemental Figure 4). Similar to the MojV G protein, the AngV G protein lacks previously described ephrin binding residues (NiV W504, E505, T531, A532, E533, N557, and Y581) (Figure 5B and Supplemental Figure 4) (16, 47, 48).

**Figure 5.**
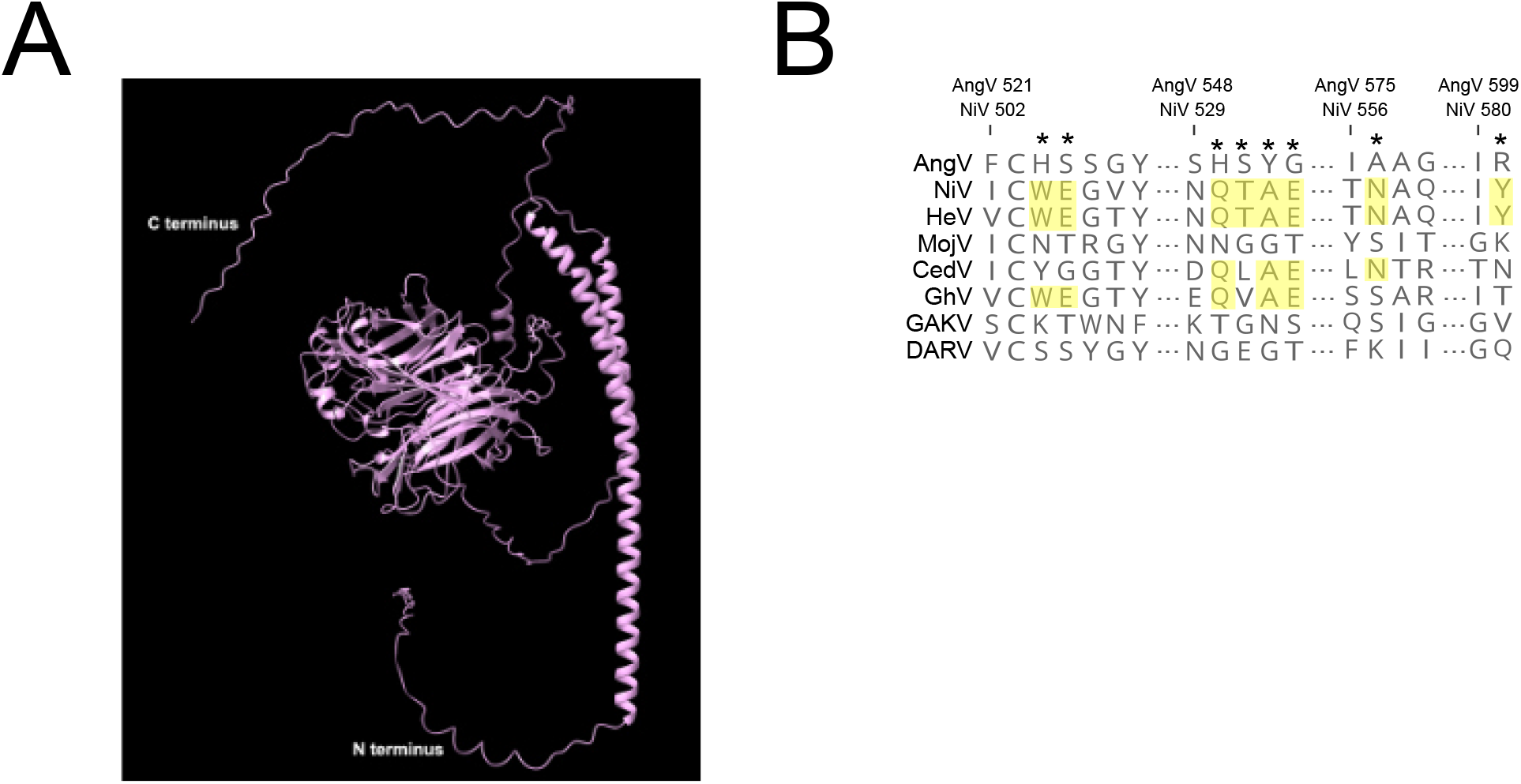
AlphaFold-predicted AngV glycoprotein 3D structure and ephrin binding residue sequence alignment. A. AlphaFold-predicted 3D structure of AngV glycoprotein. N and C termini are indicated in white. B. Alignment of HNV ephrin binding residues. The position of previously-described HNV ephrin binding residues are noted by a star, and residues conserved across most HNVs are highlighted yellow. Amino acid position numbers displayed represent the position within the AngV or NiV glycoproteins. Virus name (abbreviations) followed by GenBank Accession #: Angavokely virus (AngV) ON613535; Nipah virus (NiV) AF212302; Hendra virus (HeV) AF017149; Mojiang virus (MojV) KF278639; Cedar virus (CedV) JQ001776; Ghanaian bat Henipavirus (GhV) HQ660129; Gamak virus (GAKV) MZ574407; Daeryong virus (DARV) MZ574409.

## Discussion

We describe and characterize a novel HNV, AngV, from a urine sample collected from an *E. dupreanum* Malagasy fruit bat. In this study, urine samples from 206 unique fruit bats were assessed by metagenomic sequencing, yielding an overall positive HNV detection rate of 4.9% (10/206) for all bats studied and a HNV prevalence of 9.4% (10/106) for the *E. dupreanum* hosts. Of all the HNV positive samples, only one sample yielded sufficient reads for assembly of a complete coding sequence and subsequent genomic analysis. In a 6-year collection period spanning multiple wet/dry seasons, HNV positive samples were only recovered from *E. dupreanum* bats in the Angavokely roosting site, despite prior serological evidence of HNV infection in *P. rufus* and *R. madagascariensis* bats, as well (21). HNV RNA was recovered from *E. dupreanum* in both wet and dry seasons, though higher sampling intensity throughout the wet season precludes any conclusions regarding underlying seasonal patterns in these data. Previous work in this system has suggested a seasonal increase in fruit bat seroprevalence across the winter low nutrient season, which also overlaps the gestation period for these synchronously breeding fruit bats (21). In fruit bat systems elsewhere, HNVs are also shed in urine at higher rates during the nutrient-poor dry seasons for the localities in question (49–51); in the case of NiV and HeV, these seasonal viral shedding pulses have been linked to zoonotic spillover.

The recovered genome of AngV exhibits a structural organization characteristic of the *Henipavirus* genus and a nucleotide and amino acid identity to HeV and NiV that is comparable to those shared with the more distantly related HNVs, MojV, GhV and CedV. A limited quantity of available original sample precluded full genome recovery for AngV (as evidenced by the lack of the 5’ UTR region of the N ORF), which prevented analysis of the extent to which the full AngV genome may abide by the ‘Rule-of-Six’, observed by all other members of the *Orthoparamyxovirinae* subfamily (52). Phylogenetic analyses of AngV support classification of this virus as a distinct novel bat-borne *Henipavirus* (L gene amino acid distance <0.82 distance for the subfamily *Orthoparamyxovirinae*), in accordance to the International Committee on Taxonomy of Viruses (ICTV) criteria (19). This novel HNV is estimated to have diverged approximately 9,800 years ago, prior to the currently known African and Asian bat-borne HNV lineages but considerably more recently than the estimated mid-to late-Miocene divergence of *E. dupreanum* from its sister species, *E. helvum*, on the African continent (53). Recent characterization of *Betacoronaviruses* in Madagascar fruit bats demonstrates surprising identity to lineages circulating in West Africa (22), suggesting that, despite their endemism, Malagasy fruit bats likely experience some form of contact with the African continent. Of the 49 bat species that inhabit the island nation of Madagascar, nine species are widely distributed across Africa, Asia, and/or Europe, presenting opportunities for inter-species viral transmission via island-hopping. Intensified viral sampling of Madagascar’s insectivorous bat populations for HNVs thus represents an important future research priority.

As an ancestral bat-borne HNV, AngV may provide important insight into HNV evolution and pathogenesis. Similar to other paramyxoviruses, the encoded AngV P gene is able to produce multiple immunomodulatory protein products (54). One such protein product is the V protein, thought to be involved in immune evasion and considered a significant determinant of viral pathogenicity and lethality (55, 56). AngV harbors the highly conserved mRNA editing site and a predicted ORF that encodes a V protein with a conserved cysteine-rich C-terminus, suggesting that AngV has the capacity to produce a functional V protein. With the exception of the newly discovered HNVs in shrews, GAKV and DARV, all HNVs harboring a V protein have previously demonstrated evidence of human infection, highlighting the potential for AngV to cause productive infection in humans (1, 9, 12). Further studies are needed to ascertain the virulence potential and host breadth of this novel virus.

Characterization of the AngV glycoprotein (G) through AlphaFold modeling and structure-based alignments revealed a similar structural organization to other HNV glycoproteins. Notably, the AngV glycoprotein surpasses that of GhV as the longest glycoprotein of the *Henipavirus* genus. Like that of GhV, the AngV glycoprotein harbors a long C terminal extension (Supplemental Figure 4). It is unclear if the C terminal extension of the AngV glycoprotein has a functional role, though the C terminal extension of the glycoprotein in GhV is known to play a functional role in receptor-mediated fusion (15).

Henipavirus host tropism and virulence rely on a myriad of factors, one of which is the HNV glycoprotein. The previously characterized HNV glycoproteins of NiV, HeV, CedV, and GhV, utilize members of the ephrinA and ephrinB class family as host-cell receptors for viral entry into human cells (15–17, 47, 57). However, like MojV, the AngV glycoprotein lacks these well-conserved ephrin binding residues. Structure-based alignments can shed light on potential receptor binding residues when characterizing novel viruses. For instance, sequence-based comparisons of the GhV and NiV glycoproteins were used to predict GhV ephrin binding (12), which was later confirmed by crystallography (15). Structure-based alignment of the AngV glycoprotein shows a lack of highly conserved ephrin binding residues, including NiV E533 – a seminal residue for ephrinB2 binding that is conserved across all ephrin binding HNVs. This suggests that, like MojV—and probably DARV and GAKV—the AngV glycoprotein may not bind ephrins, pointing to the possible use of an ancestral viral entry pathway. The growing number of novel HNVs that appear not to rely on ephrin binding for cellular entry could warrant re-evaluation of the existing HNV genus to better reflect conserved function and pathobiology.

This work presents a novel bat-HNV, AngV, identified from a Malagasy fruit bat. AngV joins a growing group of ancestral HNVs with unknown cell-entry receptors. Discovery of the cell surface receptor for AngV represents an important future research priority that will shed light on the breadth of host range for this virus, including its zoonotic potential.

## Acknowledgements

The authors thank Anecia Gentles and Kimberly Rivera for help in the field and the lab and acknowledge the Virology Unit at the Institut Pasteur de Madagascar and Maira Phelps of the Chan Zuckerberg Biohub (CZB) for logistical support. They additionally thank Angela Detweiler, Michelle Tan, and Norma Neff of the CZB genomics platform for mNGS support. Molecular graphics and analyses were performed with UCSF ChimeraX, developed by the Resource for Biocomputing, Visualization, and Informatics at the University of California, San Francisco, with support from the National Institutes of Health R01-GM129325 and the Office of Cyber Infrastructure and Computational Biology, National Institute of Allergy and Infectious Diseases.

## Data Availability

Raw and assembled sequencing data are deposited in NCBI Bioproject PRJNA837298. The full genome of AngV is available in GenBank under Accession # ON613535. All raw data and code for figures can be obtained in our open-access GitHub repository: https://github.com/brooklabteam/angavokely-virus

## References

1. Eaton BT, Broder CC, Middleton D, Wang L-F. 2006. Hendra and Nipah viruses: Different and dangerous. Nat Rev Microbiol 4:23–35.

2. Sharma V, Kaushik S, Kumar R, Yadav JP, Kaushik S. 2019. Emerging trends of Nipah virus: A review. Reviews in Medical Virology 29.

3. Arunkumar G, Chandni R, Mourya DT, Singh SK, Sadanandan R, Sudan P, Bhargava B. 2019. Outbreak investigation of Nipah Virus Disease in Kerala, India, 2018. Journal of Infectious Diseases 219:1867–1878.

4. Murray K, Selleck P, Hooper P, Hyatt a, Gould a, Gleeson L, Westbury H, Hiley L, Selvey L, Rodwell B. 1995. A morbillivirus that caused fatal disease in horses and humans. Science 268:94–7.

5. Hsu VP, Hossain MJ, Parashar UD, Ali MM, Ksiazek TG, Kuzmin I, Niezgoda M, Rupprecht C, Bresee J, Breiman RF. 2004. Nipah virus encephalitis reemergence, Bangladesh. Emerging Infectious Diseases 10:2082–2087.

6. Gurley ES, Montgomery JM, Hossain MJ, Bell M, Azad AK, Islam MR, Molla MAR, Carroll DS, Ksiazek TG, Rota PA, Lowe L, Comer JA, Rollin P, Czub M, Grolla A, Feldmann H, Woodward JL, Breiman RF. 2007. Person-to-Person transmission of Nipah virus in a Bangladeshi community. Emerging Infectious Diseases 13:1031–1037.

7. Luby SP, Gurley ES. 2012. Epidemiology of Henipavirus Disease in Humans, p. 25–40. In Lee, B, Rota, PA (eds.), Henipavirus. Springer Berlin Heidelberg, Berlin, Heidelberg.

8. Marsh GA, de Jong C, Barr JA, Tachedjian M, Smith C, Middleton D, Yu M, Todd S, Foord AJ, Haring V, Payne J, Robinson R, Broz I, Crameri G, Field HE, Wang L-F. 2012. Cedar virus: a novel Henipavirus isolated from Australian bats. PLoS Pathogens 8:e1002836.

9. Wu Z, Yang L, Yang F, Ren X, Jiang J, Dong J, Sun L, Zhu Y, Zhou H, Jin Q. 2014. Novel Henipa-like virus, Mojiang paramyxovirus, in rats, China, 2012. Emerging Infectious Diseases 20:1064–1066.

10. Lee SH, Kim K, Kim J, No JS, Park K, Budhathoki S, Lee SH, Lee J, Cho SH, Cho S, Lee GY, Hwang J, Kim HC, Klein TA, Uhm CS, Kim WK, Song JW. 2021. Discovery and genetic characterization of novel paramyxoviruses related to the genus henipavirus in crocidura species in the republic of Korea. Viruses 13.

11. Drexler JF, Corman VM, Müller MA, Maganga GD, Vallo P, Binger T, Gloza-Rausch F, Rasche A, Yordanov S, Seebens A, Oppong S, Adu Sarkodie Y, Pongombo C, Lukashev AN, Schmidt-Chanasit J, Stöcker A, Carneiro AJB, Erbar S, Maisner A, Fronhoffs F, Buettner R, Kalko EK v, Kruppa T, Franke CR, Kallies R, Yandoko ERN, Herrler G, Reusken C, Hassanin A, Krüger DH, Matthee S, Ulrich RG, Leroy EM, Drosten C. 2012. Bats host major mammalian paramyxoviruses. Nat Commun 3:796.

12. Pernet O, Schneider BS, Beaty SM, LeBreton M, Yun TE, Park A, Zachariah TT, Bowden T a., Hitchens P, Ramirez CM, Daszak P, Mazet J, Freiberg AN, Wolfe ND, Lee B. 2014. Evidence for henipavirus spillover into human populations in Africa. Nature Communications 5:5342.

13. Amaya M, Broder CC. 2020. Vaccines to emerging viruses: Nipah and Hendra. Annual Review of Virology 7:447–473.

14. Cheliout Da Silva S, Yan L, Dang H v., Xu K, Epstein JH, Veesler D, Broder CC. 2021. Functional analysis of the fusion and attachment glycoproteins of mojiang henipavirus. Viruses 13.

15. Lee B, Pernet O, Ahmed A a., Zeltina A, Beaty SM, Bowden T a. 2015. Molecular recognition of human ephrinB2 cell surface receptor by an emergent African henipavirus. Proceedings of the National Academy of Sciences 201501690.

16. Laing ED, Navaratnarajah CK, Cheliout S, Silva D, Petzing SR, Xu Y, Broder CC, Xu K. 2019. Structural and functional analyses reveal promiscuous and species specific use of ephrin receptors by Cedar virus. Proceedings of the National Academy of Sciences 116:20707–20715.

17. Rissanen I, Ahmed AA, Azarm K, Beaty S, Hong P, Nambulli S, Duprex WP, Lee B, Bowden TA. 2017. Idiosyncratic Mòjiang virus attachment glycoprotein directs a host-cell entry pathway distinct from genetically related henipaviruses. Nature Communications 8.

18. Flick R, Walpita P, Czub M. 2006. Nipah and Hendra viral infections, p. 586–589. In Tropical Infectious Diseases. Churchill Livingstone.

19. Rima B, Buschmann AB-, Dundon WG, Duprex P, Easton A, Fouchier R, Kurath G, Lamb R, Lee B, Rota P, Wang L, Consortium IR. 2019. ICTV Virus Taxonomy Profile: Paramyxoviridae. Journal of General Virology 100:1593–1594.

20. Calain P, Roux L. 1993. The rule of six, a basic feature for efficient replication of Sendai virus defective interfering RNA. Journal of Virology 67:4822–4830.

21. Brook CE, Ranaivoson HC, Broder CC, Cunningham AA, Héraud J-M, Peel AJ, Gibson L, Wood JLN, Metcalf CJ, Dobson AP. 2019. Disentangling serology to elucidate henipa- and filovirus transmission in Madagascar fruit bats. Journal of Animal Ecology 88:1001–1016.

22. Kettenburg G, Kistler A, Ranaivoson HC, Ahyong V, Andrianiaina A, Andry S, DeRisi JL, Gentles A, Raharinosy V, Randriambolamanantsoa TH, Ravelomanantsoa NAF, Tato CM, Dussart P, Heraud J-M, Brook CE. 2022. Full genome Nobecovirus sequences from Malagasy fruit bats define a unique evolutionary history for this coronavirus clade. Frontiers in Public Health 10.

23. Brook CE, Ranaivoson HC, Andriafidison D, Ralisata M, Razafimanahaka J, Héraud J, Dobson AP, Metcalf CJ. 2019. Population trends for two Malagasy fruit bats. Biological Conservation 234:165–171.

24. Ranaivoson HC, Héraud J-M, Goethert HK, Telford III SR, Rabetafika L, Brook CE. 2019. Babesial infection in the Madagascan flying fox, Pteropus rufus É. Geoffroy, 1803. Parasites & Vectors 1–13.

25. Brook CE, Bai Y, Dobson AP, Osikowicz LM, Ranaivoson HC, Zhu Q, Kosoy MY, Dittmar K. 2015. Bartonella spp. in Fruit Bats and Blood-Feeding Ectoparasites in Madagascar. PLoS Neglected Tropical Diseases 9.

26. Kalantar KL, Carvalho T, de Bourcy CFA, Dimitrov B, Dingle G, Egger R, Han J, Holmes OB, Juan YF, King R, Kislyuk A, Lin MF, Mariano M, Morse T, Reynoso L v., Cruz DR, Sheu J, Tang J, Wang J, Zhang MA, Zhong E, Ahyong V, Lay S, Chea S, Bohl JA, Manning JE, Tato CM, DeRisi JL. 2021. IDseq-An open source cloud-based pipeline and analysis service for metagenomic pathogen detection and monitoring. Gigascience 9:1–14.

27. Bolger AM, Lohse M, Usadel B. 2014. Trimmomatic: A flexible trimmer for Illumina sequence data. Bioinformatics 30:2114–2120.

28. Dobin A, Davis CA, Schlesinger F, Drenkow J, Zaleski C, Jha S, Batut P, Chaisson M, Gingeras TR. 2013. STAR: Ultrafast universal RNA-seq aligner. Bioinformatics 29:15–21.

29. Langmead B, Salzberg SL. 2012. Fast gapped-read alignment with Bowtie 2. Nature 9:357–360.

30. Zhao Y, Tang H, Ye Y. 2012. RAPSearch2: A fast and memory-efficient protein similarity search tool for next-generation sequencing data. Bioinformatics 28:125–126.

31. Wu TD, Nacu S. 2010. Fast and SNP-tolerant detection of complex variants and splicing in short reads. Bioinformatics 26:873–881.

32. Bankevich A, Nurk S, Antipov D, Gurevich AA, Dvorkin M, Kulikov AS, Lesin VM, Nikolenko SI, Pham S, Prjibelski AD, Pyshkin A v., Sirotkin A v., Vyahhi N, Tesler G, Alekseyev MA, Pevzner PA. 2012. SPAdes: A new genome assembly algorithm and its applications to single-cell sequencing. Journal of Computational Biology 19:455–477.

33. Altschul SF, Gish W, Miller W, Myers EW, Lipman DJ. 1990. Basic local alignment search tool. Journal of Molecular Biology 215:403–410.

34. Edgar RC. 2004. MUSCLE: Multiple sequence alignment with high accuracy and high throughput. Nucleic Acids Research 32:1792–1797.

35. Darriba Di, Posada D, Kozlov AM, Stamatakis A, Morel B, Flouri T. 2020. ModelTest-NG: A new and scalable tool for the selection of DNA and protein evolutionary models. Molecular Biology and Evolution 37:291–294.

36. Kozlov AM, Darriba D, Flouri T, Morel B, Stamatakis A. 2019. RAxML-NG: A fast, scalable and user-friendly tool for maximum likelihood phylogenetic inference. Bioinformatics 35:4453–4455.

37. Felsenstein J. 1985. Confidence limits on phylogenies: An approach using the bootstrap. Evolution (N Y) 39:783–791.

38. Pattengale ND, Alipour M, Bininda-Emonds ORP, Moret BME, Stamatakis A. 2010. How many bootstrap replicates are necessary? Journal of Computational Biology 17:337–354.

39. Bouckaert R, Heled J, Kühnert D, Vaughan T, Wu CH, Xie D, Suchard MA, Rambaut A, Drummond AJ. 2014. BEAST 2: A software platform for Bayesian evolutionary analysis. PLoS Computational Biology 10.

40. Drummond AJ, Suchard MA, Xie D, Rambaut A. 2012. Bayesian phylogenetics with BEAUti and the BEAST 1.7. Molecular Biology and Evolution 29:1969–1973.

41. Rambaut A, Drummond AJ, Xie D, Baele G, Suchard MA. 2018. Posterior summarization in Bayesian phylogenetics using Tracer 1.7. Systematic Biology 67:901–904.

42. Drummond AJ, Rambaut A. 2007. BEAST: Bayesian evolutionary analysis by sampling trees. BMC Evolutionary Biology 7.

43. Yu G, Smith DK, Zhu H, Guan Y, Lam TTY. 2017. Ggtree: an R Package for visualization and annotation of phylogenetic trees with their covariates and other associated data. Methods in Ecology and Evolution 8:28–36.

44. Jumper J, Evans R, Pritzel A, Green T, Figurnov M, Ronneberger O, Tunyasuvunakool K, Bates R, Žídek A, Potapenko A, Bridgland A, Meyer C, Kohl SAA, Ballard AJ, Cowie A, Romera-Paredes B, Nikolov S, Jain R, Adler J, Back T, Petersen S, Reiman D, Clancy E, Zielinski M, Steinegger M, Pacholska M, Berghammer T, Bodenstein S, Silver D, Vinyals O, Senior AW, Kavukcuoglu K, Kohli P, Hassabis D. 2021. Highly accurate protein structure prediction with AlphaFold. Nature 596:583–589.

45. Pettersen EF, Goddard TD, Huang CC, Meng EC, Couch GS, Croll TI, Morris JH, Ferrin TE. 2021. UCSF ChimeraX: Structure visualization for researchers, educators, and developers. Protein Science 30:70–82.

46. Vanmechelen B, Bletsa M, Laenen L, Lopes AR, Vergote V, Beller L, Deboutte W, Korva M, Avšič Županc T, Goüy de Bellocq J, Gryseels S, Leirs H, Lemey P, Vrancken B, Maes P. 2018. Discovery and genome characterization of three new Jeilongviruses, a lineage of paramyxoviruses characterized by their unique membrane proteins. BMC Genomics 19.

47. Guillaume V, Aslan H, Ainouze M, Guerbois M, Fabian Wild T, Buckland R, Langedijk JPM. 2006. Evidence of a potential receptor-binding site on the Nipah virus G protein (NiV-G): Identification of globular head residues with a role in fusion promotion and their localization on an NiV-G structural model. Journal of Virology 80:7546–7554.

48. Negrete OA, Wolf MC, Aguilar HC, Enterlein S, Wang W, Mühlberger E, Su S v., Bertolotti-Ciarlet A, Flick R, Lee B. 2006. Two key residues in EphrinB3 are critical for its use as an alternative receptor for Nipah virus. PLoS Pathogens 2:0078–0086.

49. Field H, Jordan D, Edson D, Morris S, Melville D, Parry-Jones K, Broos A, Divljan A, McMichael L, Davis R, Kung N, Kirkland P, Smith C. 2015. Spatiotemporal Aspects of Hendra Virus Infection in Pteropid Bats (Flying-Foxes) in Eastern Australia. Plos One 10:e0144055.

50. Páez DJ, Giles J, Mccallum H, Field H, Jordan D, Peel AJ, Plowright RK. 2017. Conditions affecting the timing and magnitude of Hendra virus shedding across pteropodid bat populations in Australia. Epidemiology and Infection 145:3143–3153.

51. Cappelle J, Furey N, Hoem T, Ou TP, Lim T, Hul V, Heng O, Chevalier V, Dussart P, Duong V. 2021. Longitudinal monitoring in Cambodia suggests higher circulation of alpha and betacoronaviruses in juvenile and immature bats of three species. Scientific Reports 11:24145.

52. Hausmann S, Jacques J-P, Kolakofsky D. 1996. Paramyxovirus RNA editing and the requirement for hexamer genome length. RNA 2:1033–1045.

53. Shi JJ, Chan LM, Peel AJ, Lai R, Yoder AD, Goodman SM. 2014. A deep divergence time between sister species of eidolon (Pteropodidae) with evidence for widespread panmixia. Acta Chiropterologica 16:279–292.

54. Douglas J, Drummond AJ, Kingston RL. 2021. Evolutionary history of cotranscriptional editing in the paramyxoviral phosphoprotein gene. Virus Evolution 7.

55. Satterfield BA, Cross RW, Fenton KA, Agans KN, Basler CF, Geisbert TW, Mire CE. 2015. The immunomodulating v and W proteins of Nipah virus determine disease course. Nature Communications 6.

56. Patterson JB, Thomas D, Lewicki H, Billeter MA, Oldstone MBA. 2000. V and C proteins of measles virus function as virulence factors in vivo. Virology 267:80–89.

57. Negrete OA, Chu D, Aguilar HC, Lee B. 2007. Single amino acid changes in the Nipah and Hendra virus attachment glycoproteins distinguish ephrinB2 from ephrinB3 usage. Journal of Virology 81:10804–10814.

