## Supplemental Table 1, Supplemental Table 2, Supplemental Figure 1, Supplemental Figure 2, Supplemental Figure 3, Supplemental Figure 4 for "Discovery and Genomic Characterization of a Novel Henipavirus, Angavokely virus, from fruit bats in Madagascar"

**Supplemental Table 1.** Comparison of Intergenic Sequences and Transcriptional Start and Stop Signals of AngV and other HNV.

| Intergenic Region | Virus | Gene Stop | Junction | Gene Start |
|---|---|---|---|---|
| **N-P** | AngV | TTAGAAAAAA | CTT | AGGAGCCAAGT |
| | NiV | TTAAGAAAAA | CTT | AGGAACCAAGA |
| | HeV | TTAAGAAAAA | CTT | AGGATCCAAGA |
| | MojV | TTAAACAAAA | CTT | AGGATCCAAG |
| | CedV | TTACAAAAAA | CTT | AGGATCCAAG |
| | GhV | CTATAAAAAA | CTT | AGGATCACAA |
| | GAKV | TTAAGAAAAA | CTT | AAGAATCAAA |
| | DARV | TTATAAAAAA | CTT | AGGATGCAAG |
| **P-M** | AngV | TTATAAAAAA | CTT | AGGATCAACGA |
| | NiV | TTAAGAAAAA | CTT | AGGAGACAGGT |
| | HeV | TTAAGAAAAA | CTT | AGGAGACAGGT |
| | MojV | TCATAAAAAA | CTT | AGGAGTCAAG |
| | CedV | TTAGAAAAAA | CTT | AGGATCCCAG |
| | GhV | AGGATCACAA | CTT | AGGGATCAAG |
| | GAKV | TTAAGAAAAA | CTT | AAGAGTCAAA |
| | DARV | TTAAGAAAAA | CTT | AGGGGTAAAG |
| **M-F** | AngV | TTAAGGAAAA | CTT | AGGAGTAAAGC |
| | NiV | TTACAAAAAA | CTT | AGGAGCCAAGC |
| | HeV | TTAAGAAAAA | CTT | AGGAGCCAAGT |
| | MojV | ATATAAAAAA | CTT | AGGTGTCAGG |
| | CedV | TTAAGAAAAA | CTT | AGGATCCCAG |
| | GhV | CTAAACAAAA | CTT | AGGAAATCAG |
| | GAKV | TTAGAAAAAA | CTT | AAGAATCAAA |
| | DARV | TTAAAGAAAA | CTT | AGGACGTCAA |
| **F-G** | AngV | TTAGAAAAAA | CTT | AGGATCCAAG |
| | NiV | TTAATAAAAAA | CTT | AGGACCCAGGT |
| | HeV | TTACAAAAAA | CTT | AGGACCCAAGT |
| | MojV | TTAATAAAAA | CTT | AGGAGTCAGG |
| | CedV | TTAAATAAAA | CTT | AGGATCCCAG |
| | GhV | AATTAAAGAA | CTT | AATAATCGAG |
| | GAKV | TTAAGAAAAA | CTT | AAGAATCAAA |
| | DARV | TTATAAAAAA | CTT | AGGGGTCAAG |
| **G-L** | AngV | TTAAGAAAAA | CTT | AGGAGTAATGT |
| | NiV | TTAAGAAAAA | CTT | AGGACCCAGGT |
| | HeV | TTAAGAAAAA | CTT | AGGACCCAAGT |
| | MojV | TTACAAAAAA | CTT | AGGATTCACG |
| | CedV | TTAAAGAAAA | CTT | AGGATCCCAG |
| | GhV | CTAAGAAAAA | CTT | AGGAATTCAG |
| | GAKV | TTAAGAAAAA | CTT | AAGAATCAAA |
| | DARV | TTAAGAAAAA | CTT | AGGTGCAATG |
| **L** | AngV | TTAAGAAAAA | CTT | |
| | NiV | TTAAGAAAAA | CTT | |
| | HeV | TTAAGAAAAA | CTT | |
| | MojV | TTAATAAAAA | CTT | |
| | CedV | TTAAAGAAAA | CTT | |
| | GhV | TTAAGAAAAA | CTT | |
| | GAKV | TTAAGAAAAA | CTT | |
| | DARV | ATATAAAAAA | CTT | |

**Supplemental Table 2**. Comparison of 5' and 3' untranslated regions of AngV and other HNVs.

| Gene | Virus | 5' untranslated | | 3' untranslated | |
|---|---|---|---|---|---|
| | | Length | % Homology | Length | % Homology |
| N | AngV | | | 262 | |
| | NiV | | | 576 | 32.5 |
| | HeV | | | 558 | 34.8 |
| | MojV | | | 154 | 36.2 |
| | CedV | | | 334 | 39.5 |
| | GhV | | | 294 | 38.7 |
| | GAKV | | | 326 | 47.0 |
| | DARV | | | 458 | 39.2 |
| P | AngV | 221 | | 56 | |
| | NiV | 94 | 33.3 | 459 | 12.8 |
| | HeV | 95 | 34.5 | 459 | 17.6 |
| | MojV | 254 | 43.7 | 134 | 43.7 |
| | CedV | 98 | 35.5 | 192 | 19.0 |
| | GhV | 236 | 42.7 | 213 | 20.5 |
| | GAKV | 132 | 35.3 | 317 | 13.9 |
| | DARV | 324 | 35.8 | 199 | 52.2 |
| M | AngV | 33 | | 23 | |
| | NiV | 89 | 26.8 | 190 | 64.0 |
| | HeV | 89 | 57.6 | 190 | 18.6 |
| | MojV | 39 | 36.4 | 445 | 44.2 |
| | CedV | 114 | 24.3 | 408 | 7.6 |
| | GhV | 243 | 11.9 | 314 | 23.9 |
| | GAKV | 49 | 54.5 | 326 | 20.0 |
| | DARV | 165 | 20.0 | 527 | 5.1 |
| F | AngV | 149 | | 30 | |
| | NiV | 273 | 36.6 | 401 | 9.2 |
| | HeV | 261 | 34.6 | 408 | 6.7 |
| | MojV | 942 | 12.7 | 163 | 28.6 |
| | CedV | 276 | 38.9 | 88 | 23.5 |
| | GhV | 391 | 22.7 | 59 | 40.9 |
| | GAKV | 844 | 18.2 | 340 | 10.6 |
| | DARV | 988 | 17.5 | 65 | 42.4 |
| G | AngV | 82 | | 223 | |
| | NiV | 222 | 25.7 | 494 | 30.8 |
| | HeV | 222 | 27.5 | 506 | 29.4 |
| | MojV | 111 | 49.4 | 533 | 32.4 |
| | CedV | 98 | 42.0 | 139 | 40.8 |
| | GhV | 201 | 26.8 | 223 | 20.7 |
| | GAKV | 134 | 39.2 | 440 | 30.8 |
| | DARV | 289 | 29.4 | 529 | 36.0 |
| L | AngV | 245 | | 65 | |
| | NiV | 142 | 38.1 | 57 | 45.5 |
| | HeV | 142 | 40.0 | 57 | 40.0 |
| | MojV | 212 | 44.6 | 40 | 35.8 |
| | CedV | 293 | 42.1 | 63 | 50.8 |
| | GhV | 576 | 29.0 | 67 | 36.9 |
| | GAKV | 355 | 41.1 | 14 | 64.3 |
| | DARV | 593 | 28.6 | 68 | 32.6 |

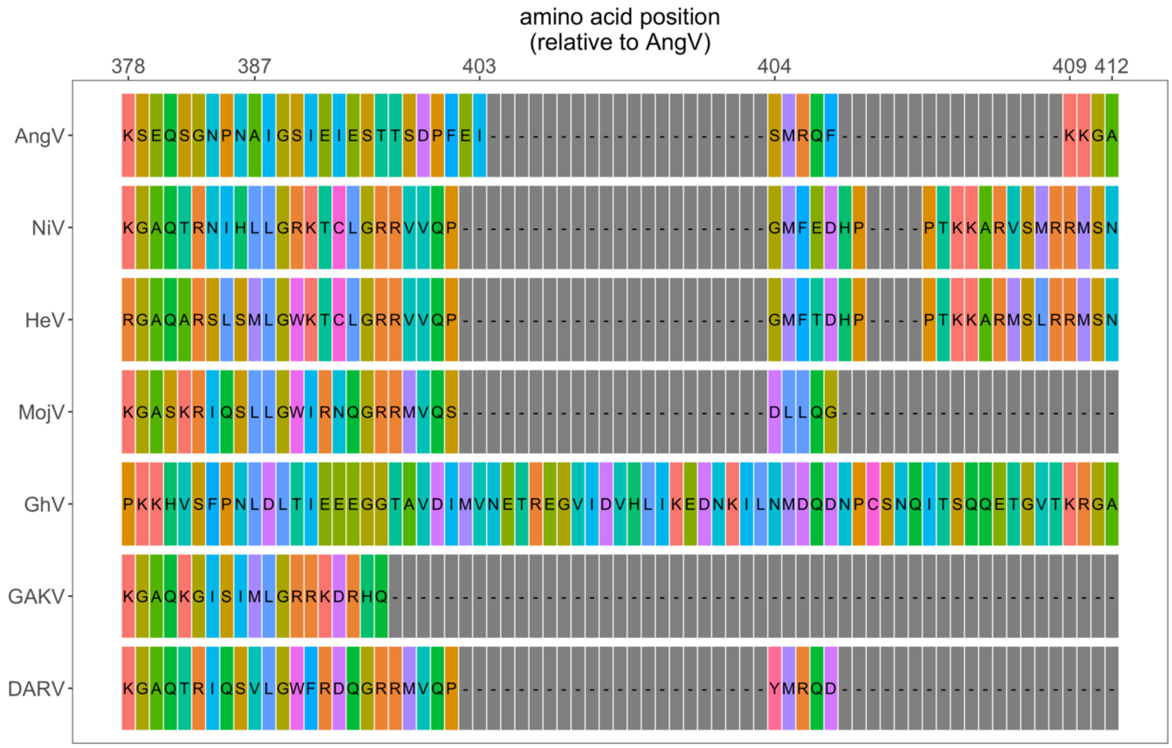


**Supplemental Figure 1.** Amino acid alignment of the C terminal region of the W protein following the addition of two guanine nucleotides to the putative mRNA editing site. The C-terminal domain of the henipavirus W protein is not conserved to the same extent as the C-terminal domain of the V protein (1). Amino acids are color coded, and position numbers represent amino acid position in the AngV protein. Virus name (abbreviation), followed by GenBank Accession #: Angavokely virus (AngV) ON613535; Nipah virus (NiV) AF212302; Hendra virus (HeV) AF017149; Mojiang virus (MojV) KF278639; Ghanaian bat Henipavirus (GhV) HQ660129; Daeryong virus (DARV) MZ574409; Gamak virus (GAKV) MZ574407. CedV was not included in panel C alignment because it does not encode a W protein.

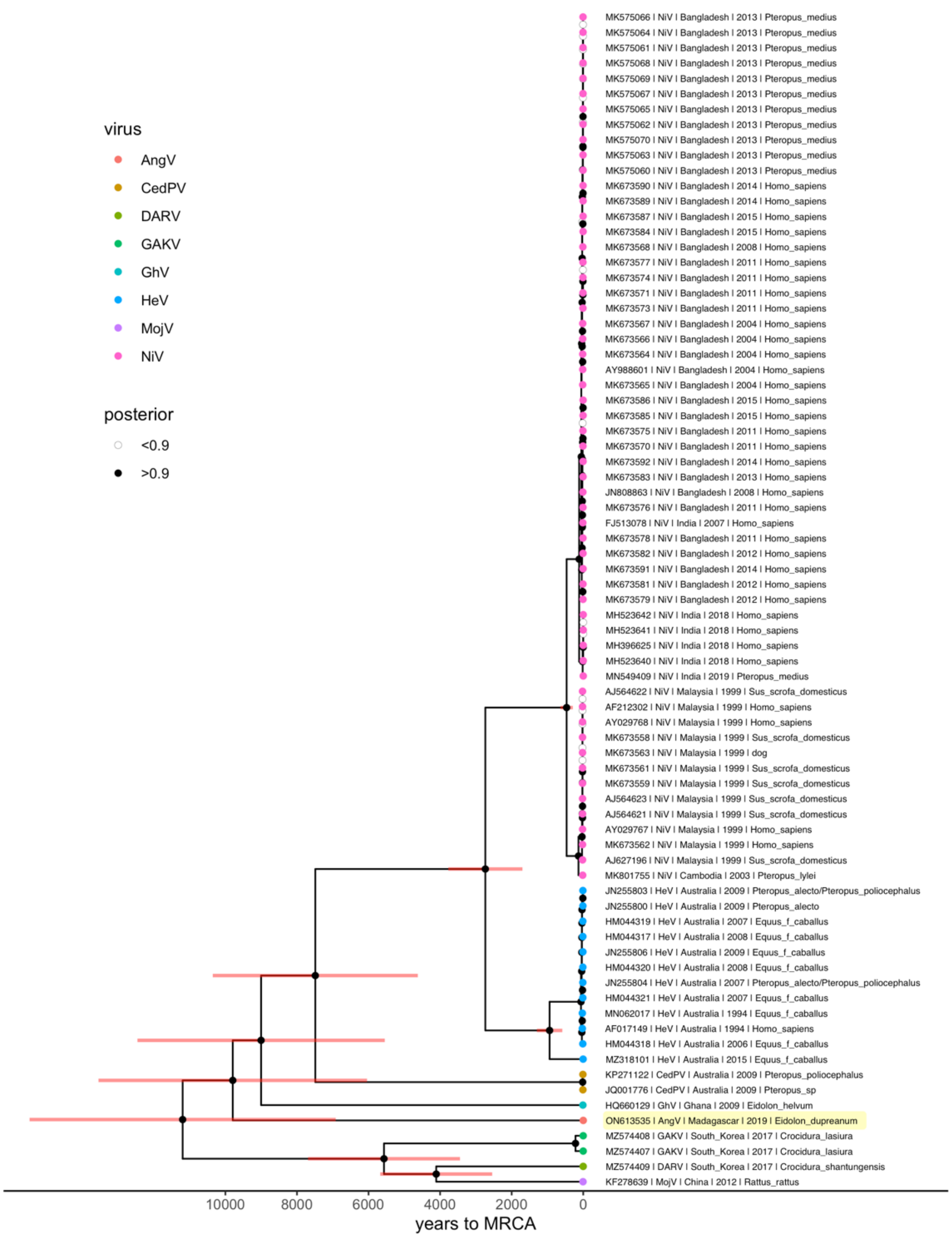


**Supplemental Figure 2.** Uncollapsed Bayesian skyline plot of all available *Henipavirus* whole genomes, with the addition of newly discovered GAKV, DARV, and AngV. Nodes with 95% credibility interval are depicted as red lines. *Henipavirus* species are highlighted by colored tip points. The estimated time to MRCA for Angavokely virus and the previously-described bat-borne HNVs is 9,794 (95% HPD: 6,519 – 14,025) years ago.

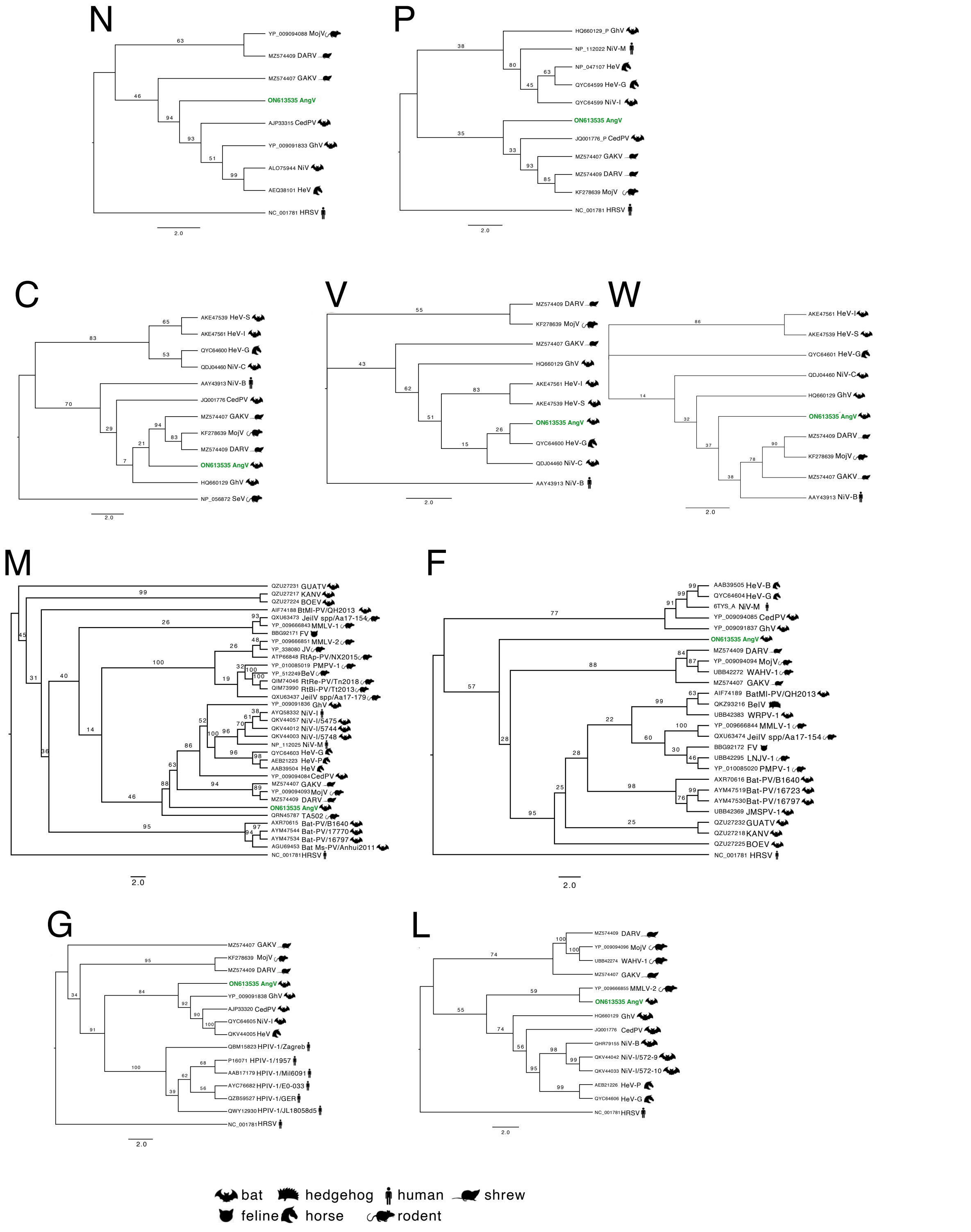


**Supplemental Figure 3.** Phylogenetic trees of AngV proteins and top BLASTx hits. Rooted phylogenetic trees, with HRSV or Sendai virus (SeV) as an outgroup. Trees for V and W proteins are unrooted. Novel HNV, AngV depicted in green. Icon represents host from which depicted virus was isolated. GenBank Accessions depicted next to viral abbreviation. Bootstrap support displayed. Scale bars represent substitutions per site. Abbreviations used as follows: Bat Ms-ParaV Bat Miniopterus schreibersii paramyxovirus; Bat-PV Bat paramyxovirus; BelV Belerina virus; BeV Beilong virus; BtMl-ParaV Bat Murina leucogaster paramyxovirus; BOEV Boe virus; CedV Cedar virus; DARV Daeryong virus; FeV Feline paramyxovirus; GAKV Gamak virus; GhV Ghana bat virus; GUATV Guato paramyxovirus; HeV Hendra virus: -B Brisbane, -I Ingham siolate, -G Gympie isolate, -Proserpine isolate, -S Sandgate Isolate; HRSV Human orthopneumovirus; JeilV spp Jeilong virus species; JMSPV-1 Jingmen Miniopterus schreibersii paramyxovirus 1; JV J virus; KANV Kanhgag paramyxovirus; MojV Mojiang virus; MMLV-1 Mount Mabu Lophuromys virus 1; MMLV-2 Mount Mabu Lophuromys virus 2; NiV Nipah virus: -C Cambodia isolate, -B Bangladesh isolate, -I India isolate, -M Mayasia isolate; PMPV-1 Pohorje Myodes paramyxovirus 1; RtAp-ParaV Rat Apodemus peninsulae Paramyxovirus; RtBi-ParaV Rat Bandicota indica paramyxovirus; RtRe-ParaV Rattus exulans paramyxovirus; SeV Sendai virus; TA502 Ruloma virus; WAHV-1 Wenzhou Apodemus agrarius henipavirus 1; WRPV-1 Wufeng Rhinolophus pearsonii paramyxovirus 1.

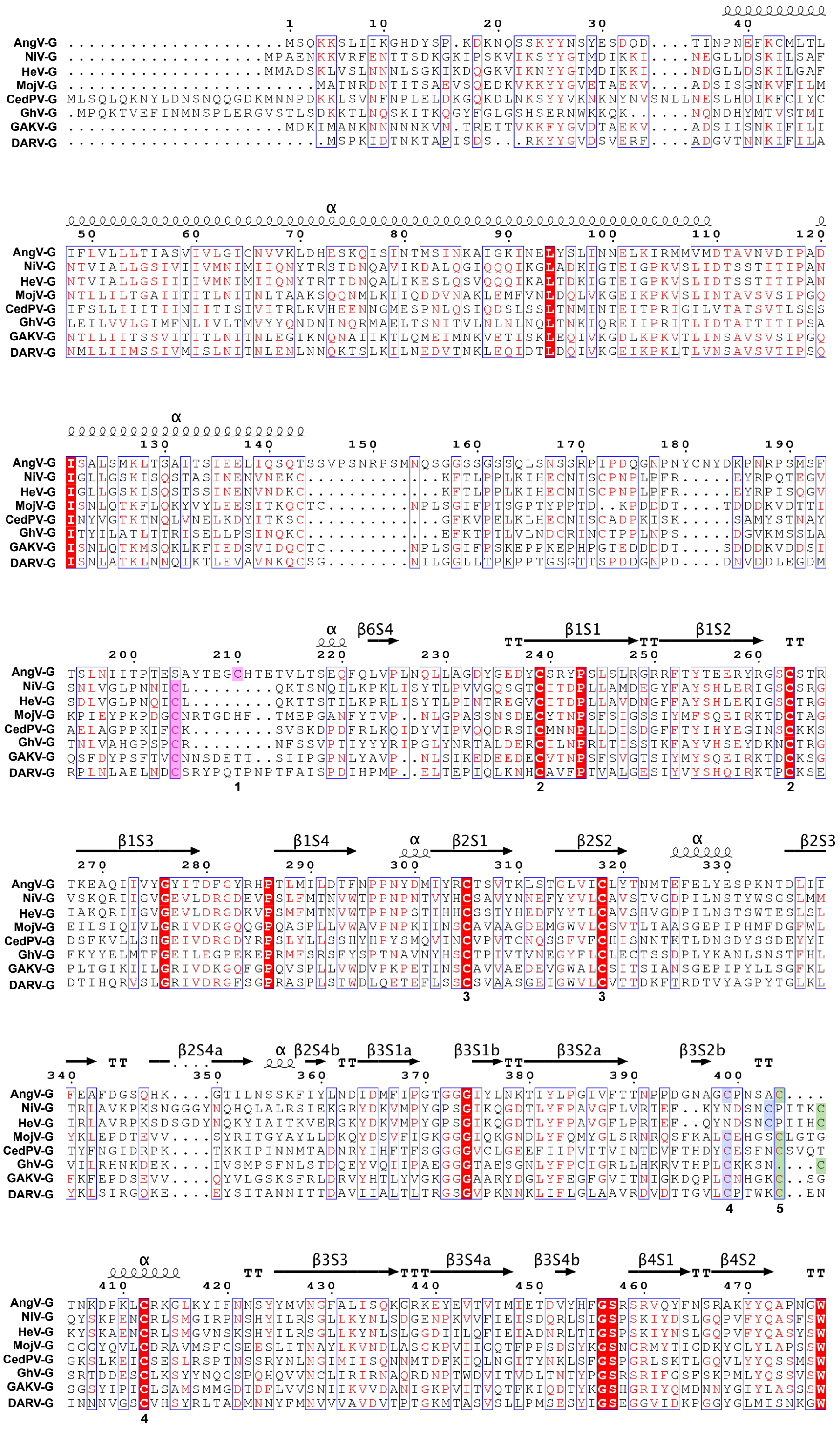
AngV-G
NiV-G
HeV-G
MojV-G
CedPV-G
GhV-G
GAKV-G
DARV-G

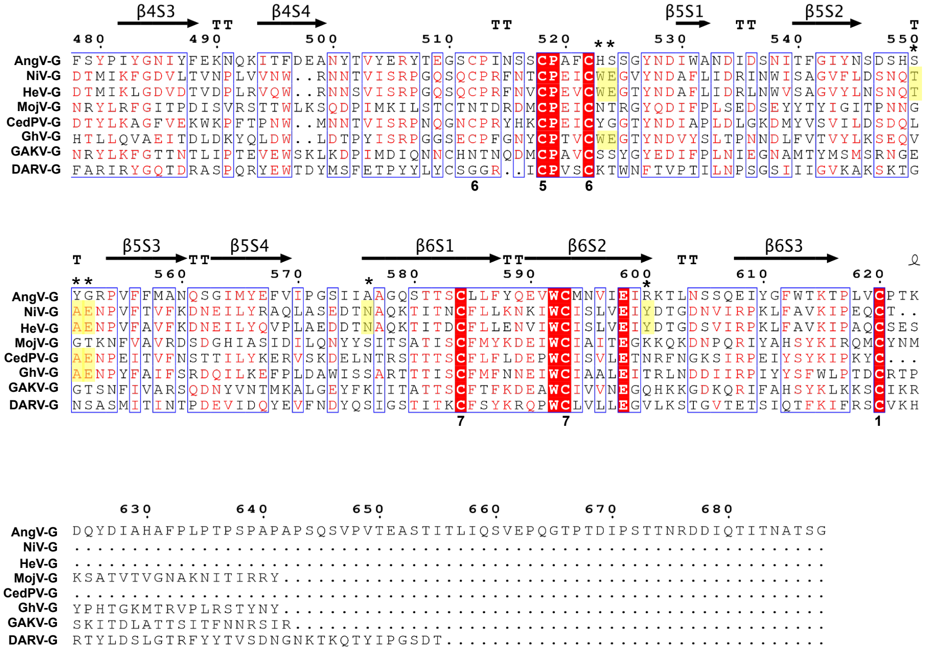


**Supplemental Figure 4.** Structure-based alignment of the HNV G protein sequences. The alignment was done with MultAlin (2) and visualized using ESPRIT (3). Residues highlighted red are fully conserved and residues colored red are partially conserved. Disulfide bonds 1 through 6 are indicated numerically underneath the alignment. Notably AngV G protein disulfide bonds 1, 4, and 5 do not align across all HNVs. Homologous residues have been highlighted pink, blue and green for disulfide bonds 1, 4, and 5, respectively. Virus name (abbreviation) GenBank Accession: Angavokely virus (AngV) ON613535, Cedar virus (CedV) JQ001776; Daeryong virus (DARV) MZ574409; Gamak virus (GAKV) MZ574407; Hendra virus (HeV) AF017149; Mojiang virus (MojV) KF278639; Nipah virus (NiV) AF212302; Ghanaian bat Henipavirus (GhV) HQ660129.

**Supplemental References**


1. Douglas J, Drummond AJ, Kingston RL. 2021. Evolutionary history of cotranscriptional editing in the paramyxoviral phosphoprotein gene. Virus Evolution 7.
2. Corpet F. 1988. Multiple sequence alignment with hierarchical clustering. Nucleic Acids Research 16:10881–108890.
3. Gouet P, Courcelle E, Stuart DI, Métoz F. 1999. ESPript: analysis of multiple sequence alignments in PostScript. Bioinformatics 15:305–308.
